# Giants among Cnidaria: large nuclear genomes and rearranged mitochondrial genomes in siphonophores

**DOI:** 10.1101/2023.05.12.540511

**Authors:** Namrata Ahuja, Xuwen Cao, Darrin T. Schultz, Natasha Picciani, Arianna Lord, Shengyuan Shao, David R. Burdick, Steven H.D. Haddock, Yuanning Li, Casey W Dunn

## Abstract

Siphonophores (Cnidaria:Hydrozoa) are abundant predators found throughout the ocean and are important components in worldwide zooplankton. They range in length from a few centimeters to tens of meters. They are gelatinous, fragile, and difficult to collect, so many aspects of the biology of these 190 species remain poorly understood. To survey siphonophore genome diversity, we performed Illumina sequencing of 32 species sampled broadly across the phylogeny. Sequencing depth was sufficient to estimate nuclear genome size from k-mer spectra in 8 specimens, ranging from 0.7-4.8Gb. In 6 specimens we got heterozygosity estimates between 0.7-5.3%. Rarefaction analyses indicate k-mer peaks can be absent with as much as 30x read coverage, suggesting minimum genome sizes range from 1.0-3.8Gb in the remaining 27 samples without k-mer peaks. This work confirms most siphonophore nuclear genomes are large, but also identifies several with reduced size that are tractable targets for future siphonophore nuclear genome assembly projects. We also assembled mitochondrial genomes for 32 specimens from these new data, indicating a conserved gene order among Hydrozoa, Cystonectae and some Physonectae, also revealing the ancestral gene organization of siphonophores. There then was extensive rearrangement of mitochondrial genomes within other physonects and in Calycophorae, including the repeated loss of atp8. Though siphonophores comprise a small fraction of cnidarian species, this survey greatly expands our understanding of cnidarian genome diversity. This study further illustrates both the importance of deep phylogenetic sampling and the utility of Illumina genome skimming in understanding genomic diversity of a clade.

**Significance:** Descriptions of basic genome features, such as nuclear genome size and mitochondrial genome sequences, remain sparse across many clades in the tree of life, leading to over generalizations from very small sample sizes and often limiting selection of optimal species for genome assembly efforts. Here we use Illumina genome skimming to assess a variety of genome features across 35 siphonophores (Cnidaria). This deep dive within a single clade identifies six species that are optimal candidates of future genomic work, and reveals greater range in nuclear genome size and diversity of mitochondrial genome orders within siphonophores than had been described across all Cnidaria.

## Introduction

Siphonophores (Figure 1) are among the longest in body length and most abundant animals in the ocean. They have a critical position in the food web of the open ocean (Hetherington et al. 2022), and many unique biological features (Mapstone 2014; Munro et al. 2018). Like corals, they are colonial animals with many highly integrated, genetically identical bodies that all arise from a single embryo by asexual reproduction. Unlike corals, they are free-swimming, and the bodies (zooids) within the colonies are specialized for different tasks, such as feeding, locomotion, and sexual production of new colonies. Studies on siphonophores have been limited, though, because they are fragile and difficult to collect. Improved genomic resources would greatly facilitate work on all aspects of their study and make it possible to learn much more from each of the specimens that are collected.

**FIG. 1.**
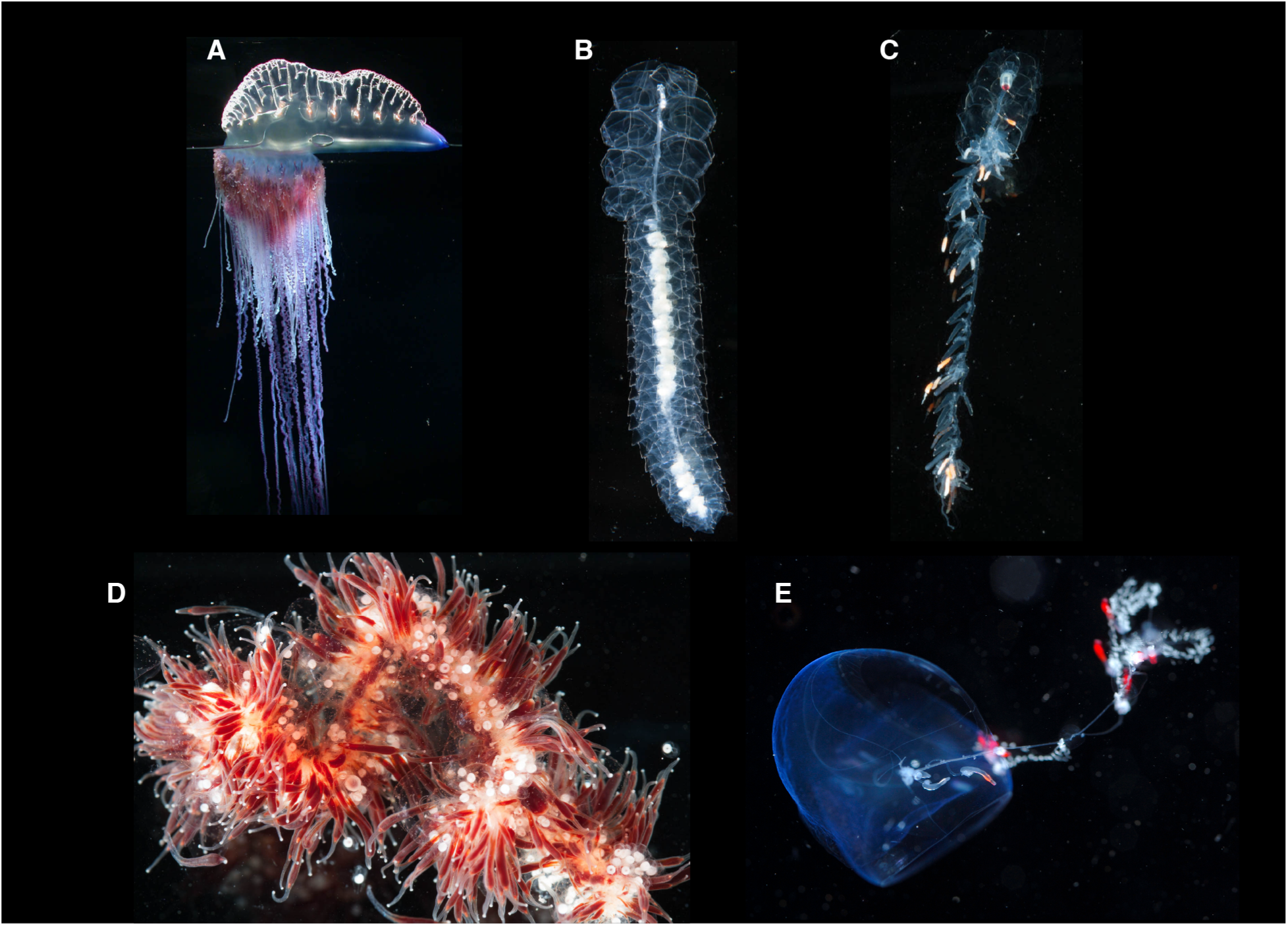
Photographs of some siphonophore species included in this study. Lengths are approximate. A. *Physalia physalis* (float is about 20mm across) B. *Frillagalma vityazi* (about 8cm long) C. *Nanomia bijuga* (about 5cm long) D. *Apolemia rubriversa* (specimen image about 7cm across) E. *Gymnopraia lapislazula* (blue nectophore is about 2cm across).

Molecular data available for siphonophores include a few genes sequenced across a large number of species (Dunn et al. 2005), transcriptomes for developmental and phylogenetic work (Siebert et al. 2011; Munro et al. 2022, 2018), and mitochondrial genomes for a couple of species (Kayal et al. 2015). Chromosome-scale genomes have been assembled for a broad diversity of cnidarians in recent years (Figure 2), but most of these are for hexacorals and relatively few are from Hydrozoa, the clade that includes Siphonophora. While no nuclear siphonophore genomes have been sequenced, two genome size estimates based on flow cytometry are available: *Agalma elegans* has been estimated to be 3.482Gb and *Physalia physalis* at 3.247 Gb (Adachi et al. 2017). These are by far the largest genomes known to exist for cnidarians, similar in size to human genomes. However, with a sample size of two out of nearly two hundred known species, sampling is insufficient to confidently describe nuclear genome size diversity within Siphonophorae.

**FIG. 2.**
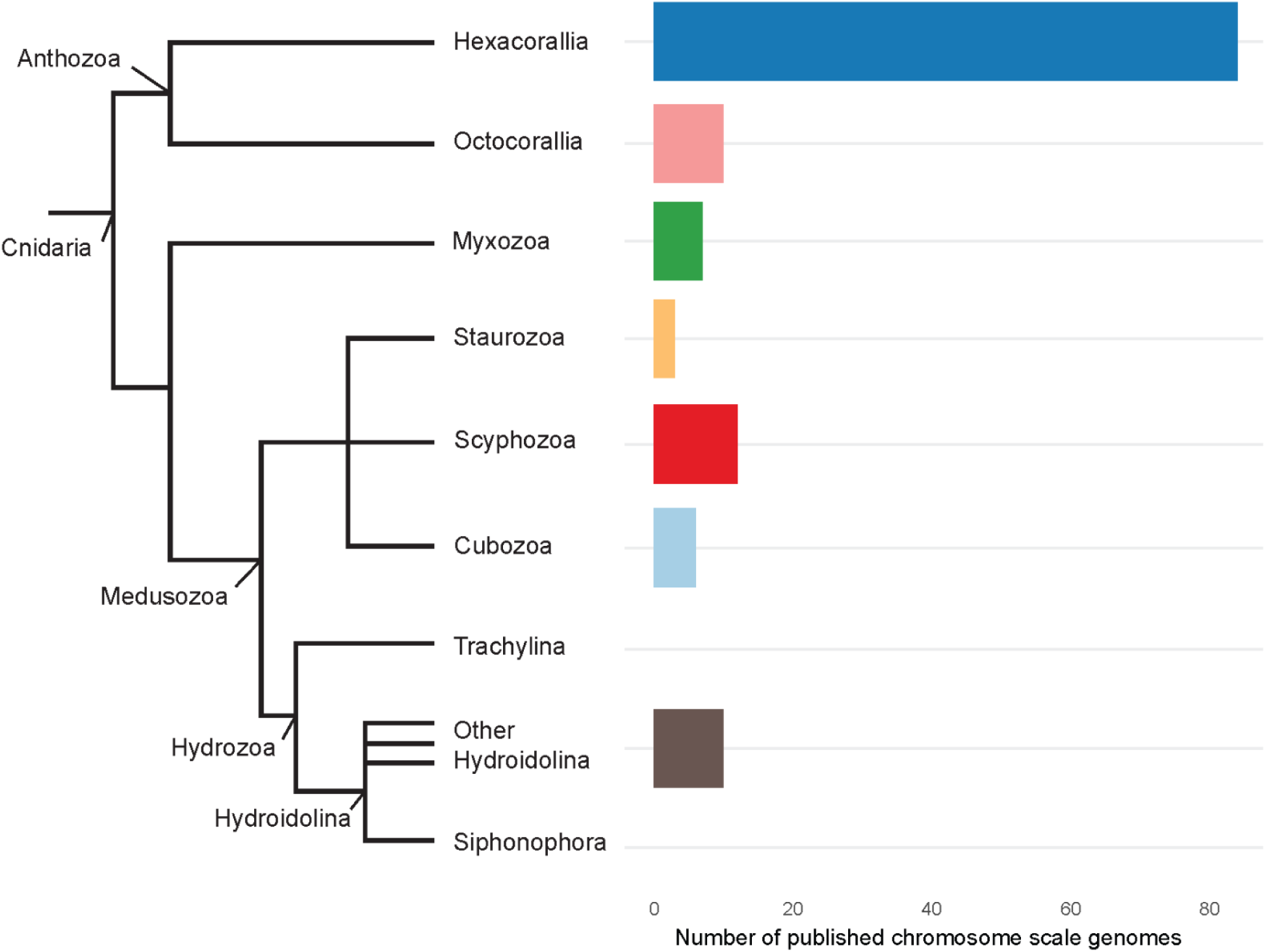
Number of chromosome-scale nuclear genomes sequenced across Cnidaria. Siphonophores are within Hydrozoa.

In parallel to nuclear genomics, considerable progress has been made in understanding cnidarian mitochondrial genomics, but this sampling is not uniformly distributed. Currently, mitochondrial genome assemblies are available for over 300 anthozoans and 70 medusozoans; however, only about 30 medusozoan mitochondrial genomes are complete. Almost all sequenced anthozoan mitochondrial genomes are circular. Though medusozoan mitochondrial genomes are thought to be linear (Bridge et al. 1992; Kayal et al. 2015), several medusozoan mitochondrial genomes in public archives are annotated as being circular. However, there has not been enough data on siphonophore mitochondrial genomes to distinguish how they may compare to other Medusozoa, especially given their semi uncertain position within the clade and the subclade Hydroidlina. Currently, there are 3 nearly-complete mitochondrial siphonophore genomes, including *Physalia physalis*, *Nanomia bijuga* and *Rhizophysa eysenhardtii (Kayal et al. 2015)*. These have identical gene orders to each other, consistent with few evolutionary changes in mitochondrial genome structure in Siphonophorae. However, the partial nature of these genome sequences and limited taxon sampling is too small to robustly test this.

Genome skimming by shotgun Illumina sequencing has emerged as a powerful tool for gaining insight about multiple genomic features (Hogan et al. 2022; Heath-Heckman & Nishiguchi 2021). Illumina reads are too short to assemble full nuclear genomes, but are sufficient to assemble mitochondrial genomes and to run a variety of analyses on nuclear genome properties based on k-mer spectra. With sufficient depth, Illumina sequencing provides estimates of nuclear genome size, heterozygosity, and repeat fraction. Here we apply genome skimming to a broad diversity of 35 specimens from 32 siphonophore species. Our objectives are primarily biological: we seek to expand sampling to understand basic features of genome diversity across siphonophores. Our objectives are also technical: we hope to identify the species that would be most amenable to chromosome-scale genome sequencing projects and evaluate the resources and approaches that would be needed for such studies at this time.

## Results

### Specimen sampling and sequencing

A total of 2,978 Gigabases (Gb) of sequence data were collected across 35 specimens from 32 species (Table 1). We took an iterative approach to sequencing, with a shallow first pass of most samples and then deeper sequencing in a progressively narrower subset of samples. The amount of data per sample therefore ranged widely, from 29 to 311Gb (Table 1). We prioritized samples according to two criteria. First, we collected more data for a specimen if we observed a peak but needed more data for the GenomeScope2 models to fit well. Second, we prioritized several abundant species that are likely to be the focus of future genomic work, including *Physalia* and *Agalma elegans*, even though it was clear that their genomes are quite large. The uneven sampling effort reflects that for samples with low sequencing depth, genomes may or may not be larger as compared to those samples with large amounts of data that we obtained clear genome size estimates.

**Table 1.**
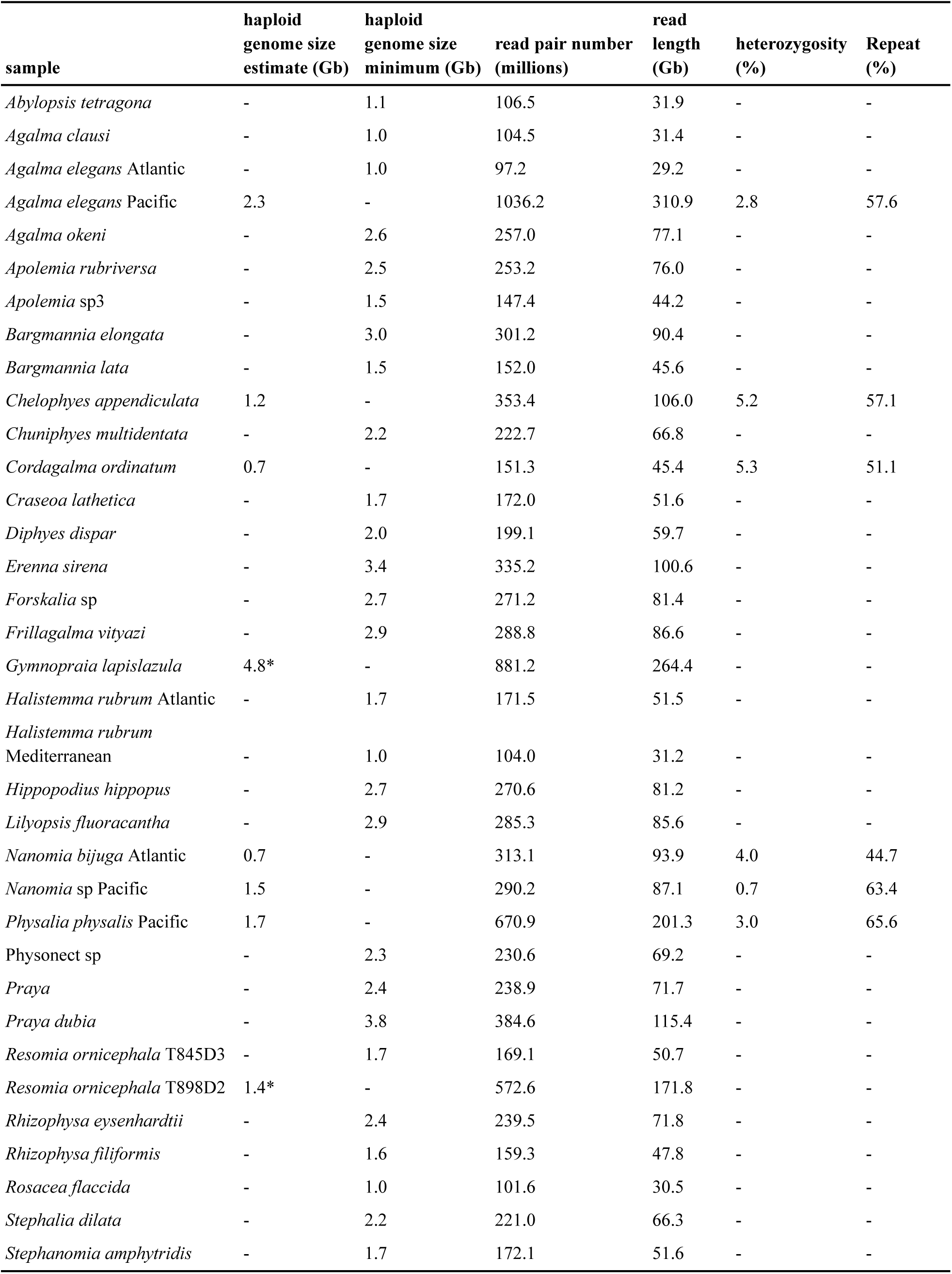
Overview of all samples. Read pair number is the number of paired 150bp reads. Read length is 150bp times the read number. Genome size minimum is provided for samples without a peak (those not in Figure 3). Given that the rarefaction analyses with as much as 30x coverage did not have a peak (Supplementary Figure 1), the genome size minimum for those samples without a peak is calculated as 1/30 the total of read length. Genome size estimate is from GenomeScope2 analyses for samples with good model fit, and from the manual estimation method for samples without good model fit (indicated by*). Heterozygosity and repeat fraction are from GenomeScope2, and are therefore only available for samples with good model fit.

### k-mer analysis

Eight samples had enough sequencing coverage to identify peaks in the k-mer spectra (Figure 3). Of these 8 samples, 6 have good model fits in GenomeScope2 (Table 1) as determined by the full model curve matching the histogram (top six plots in Figure 3). These specimens are *Cordagalma ordinatum* with an estimated haploid genome size of 700Mb, *Chelophyes appendiculata* with 1.2Gb, *Agalma elegans* (Pacific) with 2.3Gb, *Nanomia bijuga* (Atlantic) with 700Mb, *Nanomia* sp. (Pacific) with 1.5Gb and *Physalia physalis* (Pacific) with a haploid genome size estimated to be 1.7Gb. GenomeScope2 also provides estimates of heterozygosity and repeat percentage for these 6 specimens (Table 1). The 2 species that had peaks but did not display good GenomeScope2 model fit were *Gymnopraia lapislazula* and *Resomia ornicephala* (bottom 2 plots in Figure 3).

**Figure 3.**
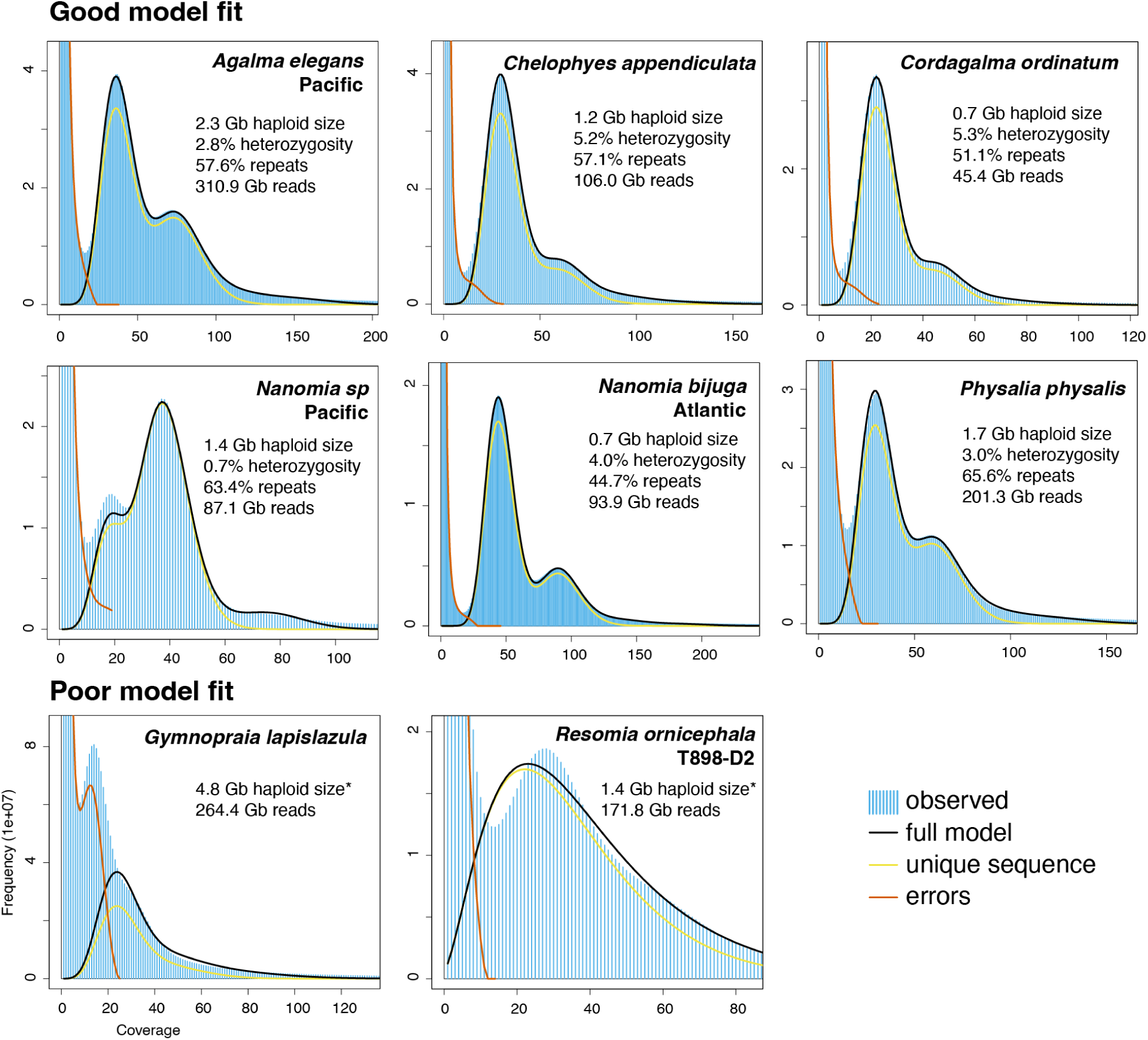
Plots of k-mer spectra for samples with peaks. Model fit is assessed by whether the full model curve (black) has peaks congruent with the underlying k-mer spectrum. The reported genome sizes for the specimens with poor model fit, indicated by asterisks, are from the manual method rather than GenomeScope2.

We also estimated genome sizes for all 8 of the species with k-mer spectrum peaks using a manual method that did not require a parametric model fit (see methods section). There is strong agreement between the estimates we obtained with this manual method and the 6 estimates obtained from the parametric GenomeScope2 models (Supplementary Table 1), suggesting that the manual method performs well. The manual method recovered haploid genome size estimates of 4.83Gb for *Gymnopraia lapislazula* and 1.40Gb for *Resomia ornicephala*.

### Rarefaction and minimal genome sizes

Sequencing was not deep enough in the other 27 sampled siphonophores to produce peaks in the k-mer spectra, so genome size estimates could not be made for these taxa. However, we can approximate a minimum genome size for each of the 27 samples based on the number of nucleotides sequenced by rarefying the data. This is done by determining the lower bound of sequencing coverage needed to produce a peak. To find the lower bound of sequence coverage needed to obtain an estimate, we first rarefied the data for three species with high confident genome estimates. (Supplementary Figure 1). Given the known genome sizes, we subsampled the reads datasets with 10x read coverage, 20x read coverage, 30x read coverage, up to 100x coverage. Because the genome sizes of each of these species is different, the number of reads in each equivalent subset was different. We then examined the k-mer spectrum for each subset: subsets with 10x and 20x read coverage did not produce peaks, two species produced peaks with 30x read coverage produced peaks, and all subsets with 40x read coverage and above produced peaks. Therefore, we conservatively estimated that samples without peaks have 30x or less read coverage. We divided the number of nucleotides sequenced by 30 to get a minimum bound on genome size. This provides minimum size estimates for the remaining genomes, ranging from 1.0-3.8Gb. These minimum estimates for samples without a peak fall entirely within the range of estimates obtained above for specimens that did have a peak. These genomes without a peak could of course be considerably larger than these minimum estimates.

Rarefaction also provides further insight into the k-mer spectra. Notably, good GenomeScope2 model fit was not achieved across all specimens until about 50x read coverage, as assessed by the full model line following the contours of the histogram (Figure 3, Supplementary Figure 1). Genome size, heterozygosity, and repeat percent estimates were generally stable with 60x and above coverage (Supplementary Figure 2).

### Mitochondrial genome structure and diversity

Mitochondrial genomes were assembled for all specimens except *Frillagalma vityazi*, *Hippopodius hippopus*, and *Lilyopsis fluoracantha*. These three unassembled taxa had 81.2-86.6Gb of data, so they were not at the low end of coverage. The issue may be that the mitochondrial genome sequences of these species differ considerably from existing databases and are not well recognized by the assembly software.

The assembler indicated that all mitochondrial genomes were linear, except that of *Chelophyes appendiculata*, which was annotated as circular. To determine whether the siphonophore mitochondrial genome is linear or circular, we split the genome sequences at *nad5* (*nad5* was chosen because it is the longest gene in the mitochondrial genome), and manually reconnected the head and tail of the original sequence, with the head of the new sequence being the second half of *nad5* and the tail being the front half of *nad5*. Sequencing reads were then mapped onto these spliced sequences. If reads spanned the head-tail junction, it indicated that the sequence is circular, otherwise it is linear. Closer inspection with read mapping revealed that the mitochondrial genome of *Chelophyes appendiculata* was linear, indicating that the circular annotation by the assembler was incorrect (Supplementary Figure 3). Nine of the 32 siphonophores mitochondrial genomes were tested using this method (Supplementary Figure 3), and they were all proved to be linear. These 9 species cover the main lineages of siphonophores, and include 2 cystonects, 3 calycophorans, 3 members of Clade A physonects, and 1 member of Clade B. These results confirm that siphonophores, like other medusozoans, have linear mitochondrial genomes.

Phylogenetic analysis of the mitochondrial protein coding genes had generally poor support for relationships within siphonophores and for the relationship of siphonophores to other hydrozoans (Supplementary Figure 4). This is consistent with low support seen for mitochondrial sequences in other groups (Kayal et al. 2015). Some nodes of the mitochondrial phylogenetic tree were constrained (Figure 4) to be consistent with major clades in previous phylogenies (Munro et al. 2018). These constraints don’t violate any nodes with 100% support in the unconstrained tree (Supplementary Figure 4), but given the poor support of the underlying tree and the use of constraints this should not be considered an independent examination of the siphonophore phylogeny.

**Figure 4.**
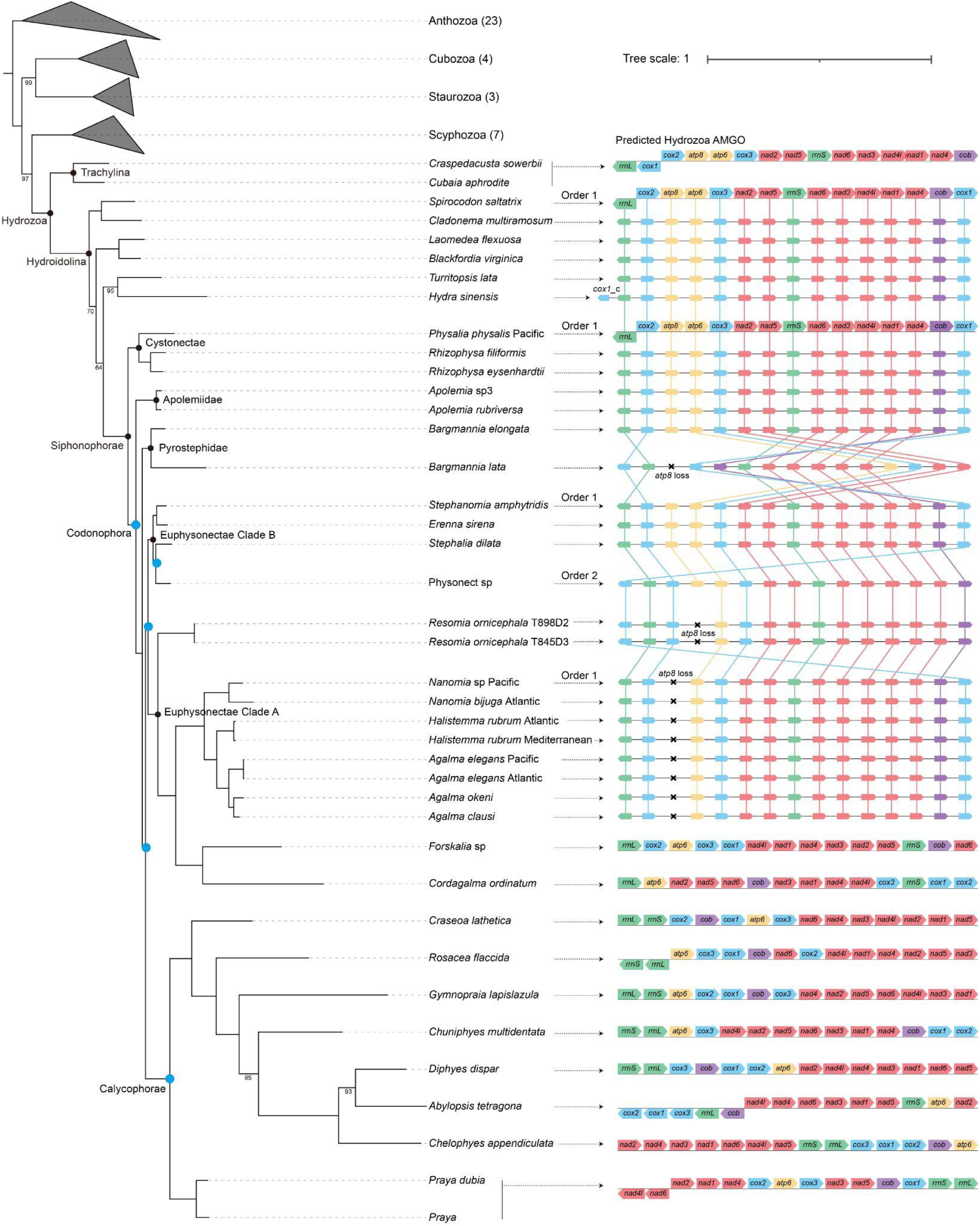
Diversity of siphonophore mitochondrial gene order. Blue circles indicate nodes constrained to be congruent with Munro et al. (2018). None of the constrained nodes had strong conflict between the mitochondrial and nuclear data, they were well supported in the nuclear data and had poor support in the mitochondrial data (see Supplementary Figure 4 for the unconstrained mitochondrial phylogeny). Bootstrap values, less than 100 are indicated at the nodes; where no numerical value is indicated at unconstrained nodes, the bootstrap value = 100. Mitochondrial rRNA and protein-coding gene arrangements of siphonophores are depicted on the right. *cox1*_c is an incomplete copy of *cox1* and lacks the 5′ end of the gene.

Mitochondrial gene order is conserved between multiple Hydrozoa taxa, Cystonectae, Apolemiidae (Fig 4), *Bargmannia elongata*, and multiple members of Clade A. This order is also found in clade B, with the modification that *atp8* has been lost. We refer to this conserved gene order as Order 1, which was also found in the two previously published cystonect mitochondrial genomes of *Physalia physalis* and *Rhizophysa* (Kayal et al. 2015).

Since Cystonectae is the sister group of Codonophora (the clade that includes all other siphonophores) and Apolemiidae is the sister group of all other codonophorans (Munro et al. 2018), this phylogenetic distribution strongly suggests that Order 1 is the ancestral mitochondrial genome order in siphonophores. *Bargmannia lata*, Physonect sp, and *Resomia ornicephala* show relatively minor variations on Order 1. However, *Forskalia* sp., *Cordagalma ordinatum*, and all members of Calycophorae show radically rearranged gene orders (Figure 4). In addition, *Bargmannia lata*, Calycophorae and Euphysonectae Clade A do not have the *atp8* gene.

Most siphonophore mitochondrial genomes we assembled have two tRNAs, one each for methionine and tryptophan. This is consistent with the tRNA inventory of many other cnidarians (Kayal et al. 2015). The exceptions were *Diphyes dispar*, *Abylopsis tetragona* and *Chelophyes appendiculata*, which had no tRNAs. We used tRNA annotation tools MITOS2 and tRNAscan-SE to verify the lack of tRNA in these species. These three species form a clade, consistent with a shared loss of tRNA in this group.

## Discussion

Our results confirm that siphonophores have the largest known genomes within Cnidaria, and reveal striking variation in genome size within the group. We were able to estimate genome sizes for eight (Table 1, Figure 3) of the 35 specimens. These specimens include some species with larger genomes that we sequenced more deeply, and others that have relatively smaller genomes for siphonophores. All, though, have very large genomes relative to other cnidarians (Figure 5). For the 27 specimens that had no k-mer spectra, we estimated their minimum genome sizes to be between 1.0-3.8Gb. These minima are quite conservative, though, and the genomes could be considerably larger.

**Figure 5.**
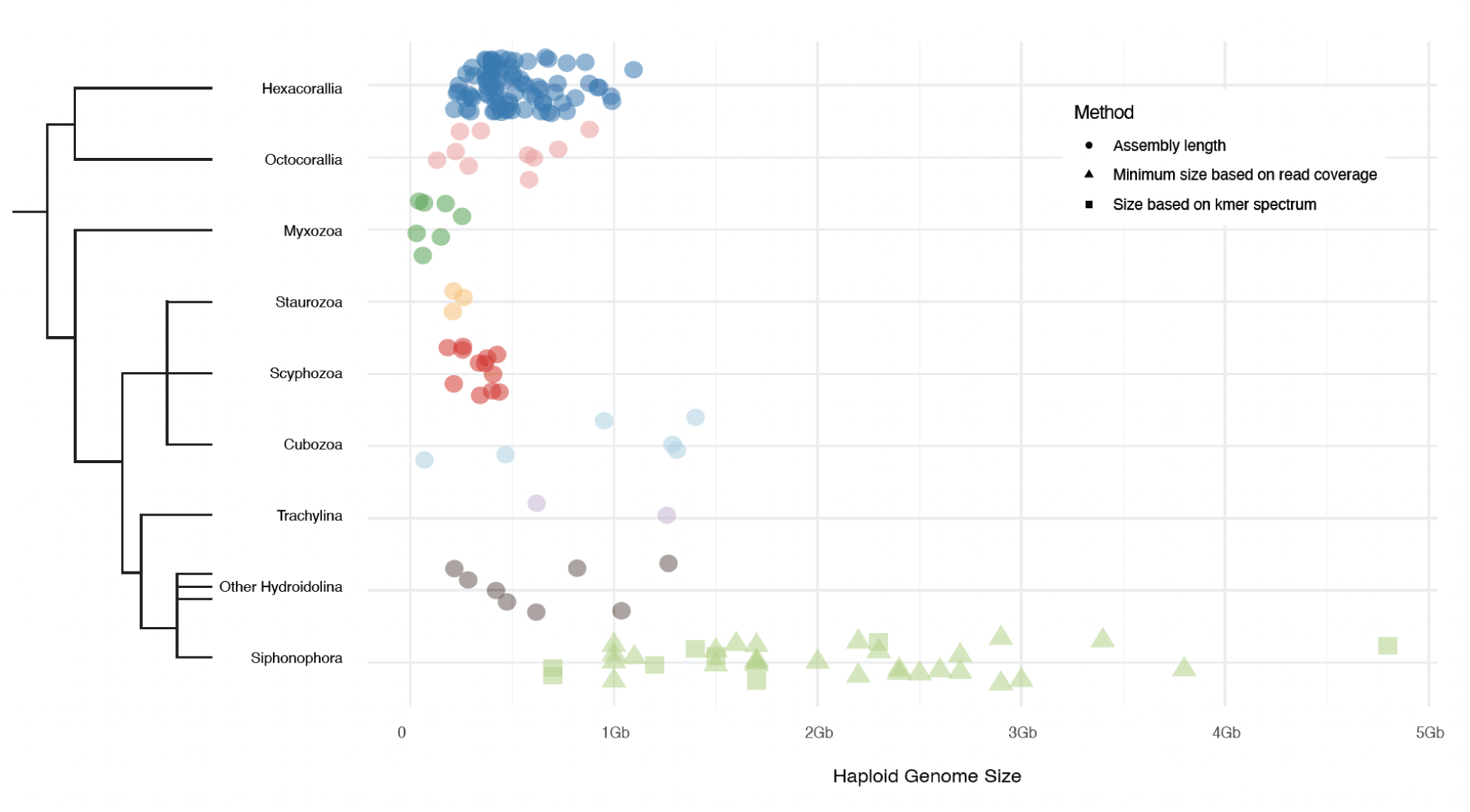
Cnidarian genome sizes and genome size estimates arranged by group. All Siphonophora represented here are newly sequenced for this project.

The smallest observed genomes are scattered across the siphonophore phylogeny and nested within groups with much larger genomes. Thus, our findings suggest that the most recent common ancestor of siphonophores already had a very large genome relative to other cnidarians, and that there have been multiple independent reductions in genome size within isolated lineages. The most striking reduction of genome size we identified was within *Nanomia*. We included two specimens, one from the Pacific and one from the Atlantic. The Pacific specimen has a genome size of 1.5Gb, within the range of many other siphonophores, while the Atlantic specimen has a genome of only 700Mb, one of the two smallest siphonophore genomes we found. Remarkably, both of these specimens come from populations that are commonly referred to as a single species, *Nanomia bijuga*, though the branch length separating them is longer than for other conspecific specimens (Dunn et al. 2005). Additionally, Haeckel(Haeckel 1888) described 8 distinct species, which Totton (British Museum (Natural History). Department of Zoology et al. 1965) later synonymized as all *Nanomia bijuga,* indicating that there are morphological differences among *Nanomia* sp. populations. Taken together, this evidence strongly suggests these are in fact different species, though a taxonomic revision is beyond the scope of this paper. In the meantime, we refer to the Atlantic specimen as *Nanomia bijuga*, and to the Pacific specimen as *Nanomia* sp. These results show that genome skimming, even without genome assembly, can provide insight into questions about systematic biology. It is notable that the genomes differ in size close to two-fold and that *Nanomia bijuga* Atlantic has a lower fraction of repeat sequences, 44.7%, than does *Nanomia* sp Pacific, 63.4% (Table 2). This indicates that a possible reduction in the size of the *Nanomia bijuga* Atlantic could be due to a reduction in repetitive regions. Full genome sequences for both of these taxa will be a fascinating opportunity to further understand radical changes in cnidarian genome size over relatively short evolutionary time scales.

Genome sizes were previously estimated for two siphonophores collected in Japan, *Agalma elegans* at 3.482Gb and *Physalia physalis* at 3.247 Gb, with cell flow cytometry (Adachi et al. 2017). These sizes differ considerably from the estimates here we obtained of 2.3Gb for *Agalma elegans* (Pacific) and 1.7Gb for *Physalia physalis* (Guam). The specimens were collected at different locations for the two studies and, as for *Nanomia*, there may be more diversity than is commonly appreciated within these species, including variation in genome sizes across populations. Alternatively, there could be technical variations between cell flow cytometry and k-mer analyses.

The 32 siphonophore mitochondrial genome assemblies revealed remarkable conservation of gene order in early siphonophore evolution, followed by bursts of changes in gene order within Eucladophora (Damian-Serrano et al. 2021). Previous studies have identified 4 main mitochondrial gene orders in Medusozoa (Kayal et al. 2015), but we found more than 10 gene orders in siphonophores. Mitochondrial gene order 1 (Figure 4) spans the most recent common ancestor of siphonophores, and is also present in outgroup taxa, suggesting that this is the AMGO (Ancestral Mitochondrial Genome Organization) of siphonophores. The mitochondrial gene order within medusozoans is well conserved, and the predicted Medusozoa AMGO is shared by most of the Scyphozoa, Staurozoa, Cubozoa (ignoring genome fragmentation), and Hydrozoa Trachylina. But the order underwent some changes in the ancestor of Hydrozoa Hydroidolina (the clade that includes siphonophores), such as loss of two non-standard protein-coding genes *polB* and ORF314, and *cox1* was inverted and translocated (Kayal et al. 2015). And the AMGO order 1 of siphonophores is consistent with the AMGO of Hydroidolina.

There are several striking features of the diversity of mitochondrial gene order within Siphonophora, one being the multiple independent losses of *atp8*. Although this is the first loss we know of in Cnidaria, this gene is absent in mitochondrial genomes of other animals including ctenophores and sponges (Schultz et al. 2020) (Haen et al. 2007) (Gissi et al. 2008). In some parasitic algae it has been lost entirely (Hancock et al. 2010). We also found that all tRNAs are lost in three species that form a clade in Calycophorae, *Diphyes dispar, Abylopsis tetragona* and *Chelophyes appendiculata.* In other non-bilaterian species, there are documented cases of tRNA loss (Lavrov & Pett 2016), including in ctenophores (Pett et al. 2011) (Pett & Lavrov 2015), some sponge lineages (Wang & Lavrov 2008) and some cnidarians (Kayal et al. 2013). The remarkable conservatism of mitochondrial gene order across Hydrozoa and many siphonophores contrasts sharply with the radical rearrangements seen within *Forskalia* sp., *Cordagalma ordinatum*, and all members of Calycophorae (Figure 4). They are so rearranged that it is difficult to find any pattern to remark upon. With the exception of *Praya* and *Craseoa lathetica*, specimens with highly reorganized mitochondrial genomes also have exceptionally long branches in the mitochondrial protein sequence phylogeny. Rearrangements of the mitochondrial gene order in siphonophores therefore tend to be associated with an elevated rate of evolution of the gene sequences themselves.

The assembler (NOVOPlasty) indicated that our assembled mitochondrial genome of *Chelophyes appendiculata* was circular. This would be very surprising given the consistent linear mt genomes of Medusozoa (Bridge et al. 1992). Further inspection with read mapping indicated that this genome is linear (Supplementary Figure MT1), and that the circular annotation was spurious. We note that some other medusozoan genomes deposited in Genbank are annotated as circular, such as *Nemopilema nomurai* (KY454767), *Rhopilema esculentum* (KY454768), *Catostylus townsendi* (OK299144), *Chrysaora pacifica* (MN448506), *Haliclystus antarcticus* (KU947038), among others. We suspected that these were also spurious annotations. SRA data is available for *Nemopilema nomurai, Rhopilema esculentum* and *Catostylus townsendi* so we tested whether the reads for these species passed through the head-tail junction. They did not (Supplementary Figure 5), affirming our suspicions that these mitochondrial genomes are actually linear. These results show that assembler softwares seem to be biased toward circular annotations, and these results should always be critically evaluated.

In addition to the biological insight gained with these genome skimming analyses, they have technical value for prioritizing future siphonophore genome projects. Though large relative to other cnidarian genomes, siphonophore genomes for which we obtained confident genome size estimates can feasibly be sequenced and assembled to chromosome scale using standard modern approaches. Given their tractable genome sizes, phylogenetic distribution, and interesting biology, these 8 species (Figure 3), apart from *Gymnopraia lapislazula* with its nearly 5Gb genome, are promising priorities for future *de novo* siphonophore genome assembly projects.

Though the roughly 190 described siphonophores make up a tiny fraction of the more than 10,000 cnidarian species, this deep dive within this single clade greatly enriches our understanding of nuclear and mitochondrial genome diversity in Cnidaria. Other poorly known clades may have similar surprises to offer and may benefit from genome skimming to better understand their genome diversity in the context of evolution.

## Materials and Methods

### Summary of genomes across Cnidaria

Cnidarian genome sizes were retrieved in February 2023 from https://www.ncbi.nlm.nih.gov/data-hub/genome/.

### Sampling

Out of the 35 specimens examined in this study, 23 were collected in a previous study of siphonophore phylogenetics (Dunn et al. 2005). The remaining 12 specimens are new. Most were collected by deep sea submersible off the coast of California by Monterey Bay Aquarium Research Institute from the research vessel *R/V Western Flyer*. We also collected a *Physalia* in Guam that was preserved in ethanol. Specimens collected by the *R/V Western Flyer* were frozen with liquid nitrogen, homogenized by mortar and pestle, and gDNA extracted with EZNA Mollusc DNA kit (Cat. #D3373-01). The *Physalia* specimen was ground up via mortar and pestle and then gDNA extracted using the Qiagen DNeasy Blood and Tissue Kit (Cat. #69504). Samples were quantified via Nanodrop and Qubit.

### Sequencing

Library preparation and sequencing was completed by the Yale Center for Genome Analysis (YCGA). Libraries were prepared with the IDT xGen DNA Library Prep EZ Kit (Cat.# 10009821). They were then sequenced on an Illumina NovaSeq6000 instrument using a S4 flow cell in a 2×150bp paired end run. The numbers of reads and bases sequenced for each specimen are indicated in Table 1.

### Genome k-mer analysis and manual genome estimation

We developed a Snakemake (Köster & Rahmann 2012) workflow that utilized Jellyfish v2.3.0 (Marçais & Kingsford 2011) to count k-mers (k=21) and GenomeScope 2.0 (Vurture et al. 2017; Ranallo-Benavidez et al. 2020) to analyze k-mer histograms. The snakemake script and other files for this analysis are available in the github repository.

In addition to the parametric models of GenomeScope2, we also applied a standard manual method for genome size estimation from k-mer spectra with a peak. This generally follows the approach outlined at https://bioinformatics.uconn.edu/genome-size-estimation-tutorial/ . The histogram produced by Jellyfish has two columns, which we refer to as X and Y. X is the count of k-mer occurrences in the input sequences, and Y is the number of k-mers with that count. We identified two X values by manual inspection of each histogram. *X_min_* is the X position of the first minimum. All counts to the left of *X_min_* are considered to be sequencing errors and are excluded from further consideration. *X_peak_* is the X position of the first peak. We then estimated the haploid genome length *L* as:

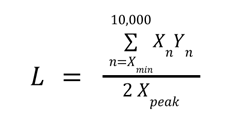

The 2 is in the denominator since the first peak is the heterozygous peak. The sum is through 10,000 since this is the maximum included in the Jellyfish histogram output.

### Rarefaction of k-mers and minimum genome sizes

To identify the read coverage (i.e., the total length of sequencing reads divided by the haploid genome size) required to observe a peak in the k-mer histogram, we subsampled the reads for *Agalma elegans* Pacific*, Cordagalma ordinata*, and *Chelophyes appendiculata*. These species were selected because they had good model fit in GenomeScope2 analyses, and therefore, good genome size estimates. We created data subsets of read coverages in increments of 10 fold coverage of haploid genome size, and then ran our same Snakemake k-mer analysis workflow on each of these subsets. Details on the number of reads included in each subset are available in Supplementary Table 4.

We did not observe peaks in data subsets with 10x or 20x read coverage. *Agalma elegans* did not have a peak at 30x coverage, but the other two rarefied specimens did. All taxa had peaks at 40x. Thus, we estimated the minimum genome size of specimens without a peak in the k-mer plot as the total length of all reads divided by 30, the highest coverage at which a taxon did not have a peak.

### Mitochondrial genome assembly and annotation

All the mitochondrial genomes were *de novo* assembled with NOVOPlasty v4.3.1 (Dierckxsens et al. 2017) except the *Bargmannia lata* mitogenome, which was assembled by GetOrganelle v1.7.6 (Jin et al. 2020). Annotation of the mitochondrial genomes was conducted with MITOS2 web server (Donath et al. 2019) with the Mold, Protozoan, and Coelenterate Mitochondrial Code and the Mycoplasma/Spiroplasma Code (NCBI translation table 4), using default settings and canceling the circular setting. Then, genome annotation of start and stop positions of each gene was manually adjusted using Artemis (Rutherford et al. 2000).

### Phylogenetic analysis

The amino acid sequences of 13 mitochondrial protein encoding genes for 32 siphonophores, 22 other medusozoans, and 23 anthozoans were used for phylogenetic analysis. See Supplementary Table 5 for accession numbers of previously published genomes included in these analyses.. Each gene was individually aligned in MAFFT v7.310 (Katoh & Standley 2013) and trimmed using Trimal v1.4.1 (Capella-Gutiérrez et al. 2009) with default parameters to discard ambiguously aligned sites. Gene alignments were then concatenated into a final supermatrix. Unconstrained maximum-likelihood (ML) phylogenetic analysis was then conducted using IQ-Tree 2.0.3 (Minh et al. 2020) with the option -m MFP and evaluated with 1,000 fast-bootstrapping replicates (Supplementary Figure MT2). Given the broader sequence sampling of the transcriptome phylogeny, we ran constrained inferences after clamping the five nodes (blue circles in Figure 4) to be consistent with the topology of the tree in Munro *et al*. (2018).

## Supplementary Material

Supplementary figures, tables and data are available in the git repository.

## Acknowledgements

Thanks to Lourdes Rojas and Eric Lazo-Wasem at the Yale Peabody Museum of Natural History. Support for this project came from multiple sources including the National Science Foundation grant NSF-OCE 1829835, the Tal Waterman Fund, and the Qingdao Postdoctoral Applied Research Project (Grant Number: QDBSH20220202163). D.T.S. was supported by ERC-H2020-EURIP grant No. 945026. Thank you to the NIH T32 Training Grant in Genetics at Yale. All library preparation and sequencing was conducted at the Yale Center for Genomic Analysis. All analyses were performed on computer clusters operated by the Yale Center for Research Computing.

## Data availability

Code and other analysis components are available at https://github.com/dunnlab/siph_skimming. Sequence reads have been deposited in NCBI SRA as BioProject PRJNA925656. Annotated mitochondrial genomes have been deposited at NCBI with accession numbers OQ957189-OQ957220.

## Author notes

NA - new sample collection, new DNA extractions, coordination with sequencing center, implemented k-mer analysis workflow, manuscript writing

DS - project design, developed k-mer analysis workflow, interpretation of k-mer spectra CWD - project design, data analysis, data submission, manuscript writing

NP - implemented k-mer analysis workflow, manuscript edits AL - summary of existing cnidarian genomes

XC - Assembly, annotation, and analysis of mitochondrial genomes, manuscript writing SHDH - new sample collection, project design

YL - Assembly, annotation, and analysis of mitochondrial genomes, manuscript writing DB - Collected *Physalia* specimen

SS - Assistance with mitochondrial genome assembly

