## Supplemental material for "Giants among Cnidaria: large nuclear genomes and rearranged mitochondrial genomes in siphonophores"

### Supplementary data

#### Supplementary files

Supplementary photographs. Images of specimens that were newly collected by remotely operated underwater vehicle (ROV). Available in the git repository.

### Supplementary figures

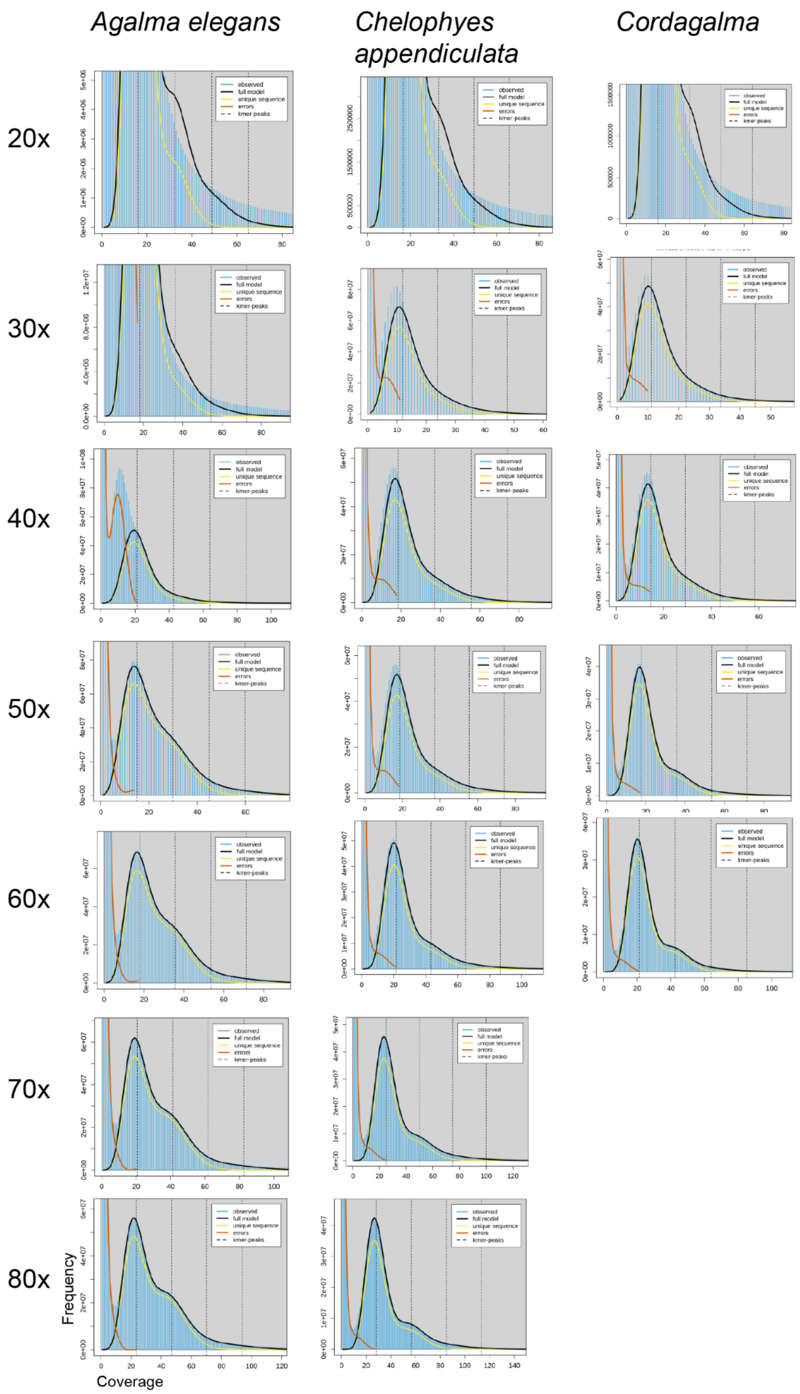


Supplementary Figure 1. Rarefaction plots. Linear plots from GenomeScope2 for three species (columns) with different read subsets (rows). The row labels indicate the read coverage of the subset relative to the haploid genome size estimated with all the data. For a species with a 1Gb genome size, for example, 20x would indicate that 20Gb of read data were included in the subset. If there were no sequencing errors and the genome were highly heterozygous, the coverage of the left (heterozygous) k-mer peak would be expected to be half the read coverage. Peaks from real data fall to the left of this expected position since many k-mers include sequencing errors (falling to the left of the minimum) or are homozygous (forming a second right peak or shoulder). Peaks are not observed with less than 20x read coverage, indicating that histograms without a peak can have 20x or less coverage. Reasonable model fit (where the full model curve follows the histogram for all three samples) requires about 50x read coverage.

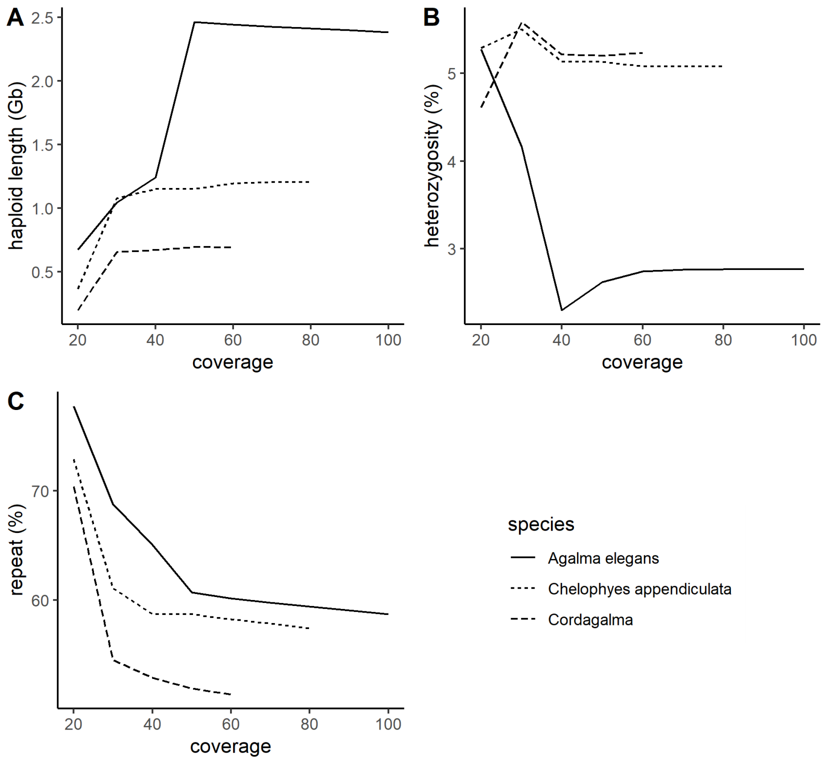


Supplementary Figure 2. The impact of increased sampling on estimates of genome length (A), heterozygosity (B), and repeats (C). Based on the same rarefaction analyses is Figure 3. In each case, the lower end of the credibility interval is presented.

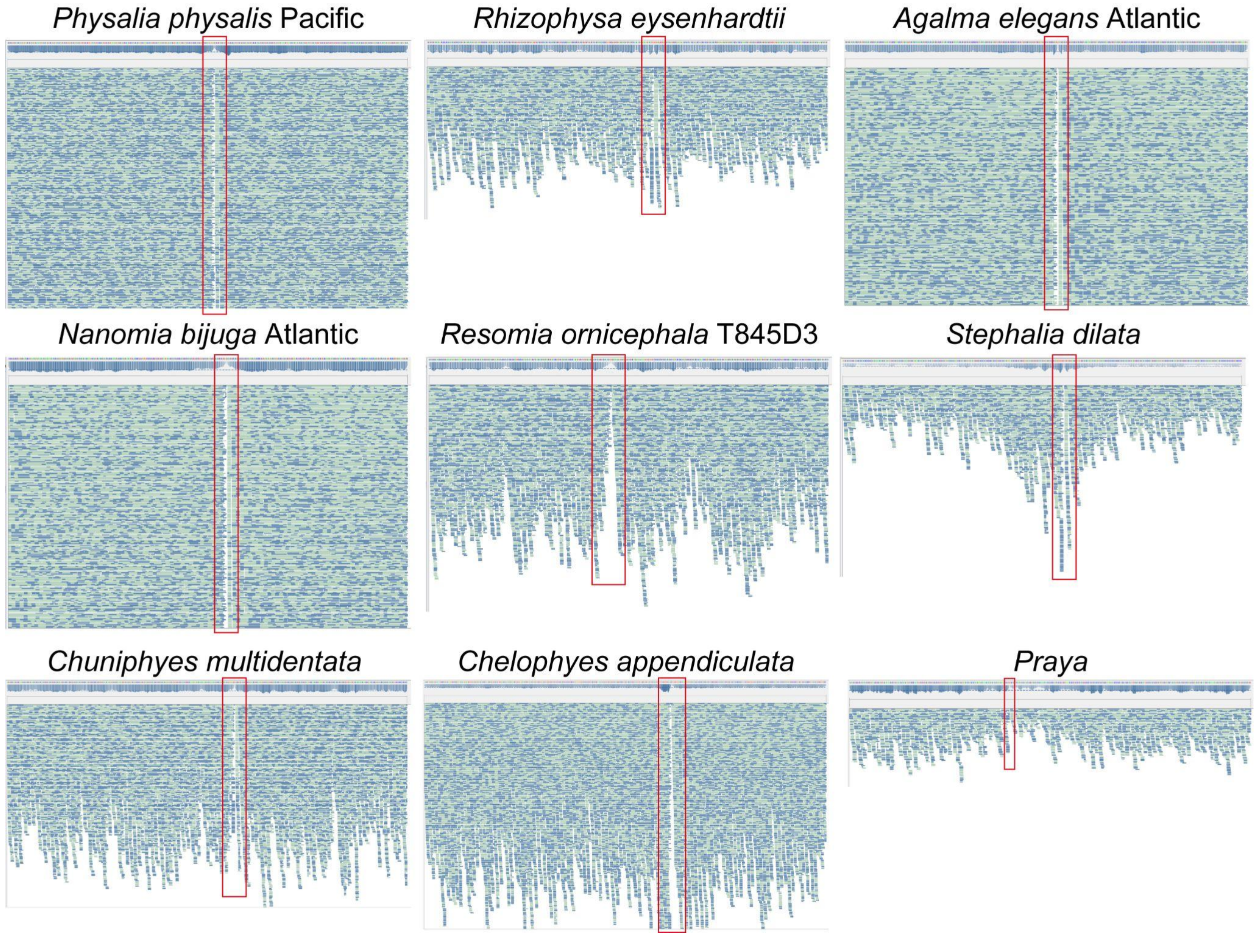


Supplementary Figure 3. Read mapping demonstrates the linear structure of siphonophore mitochondrial genomes. Mitochondrial genome sequences were split from *nad5* and concatenated head-to-tail of the original sequence, and then sequencing reads were mapped onto newly concatenated sequences. If the mitochondrial genome sequence was circular, there would be many reads passing through the head-tail junction. Our mapping results showed that there will be a gap (marked by red rectangles) at the head-tail junction of each species tested, indicating that the mitochondrial genomes of these siphonophores were linear. Green indicates the forward sequencing reads, and blue indicates the reverse sequencing reads.

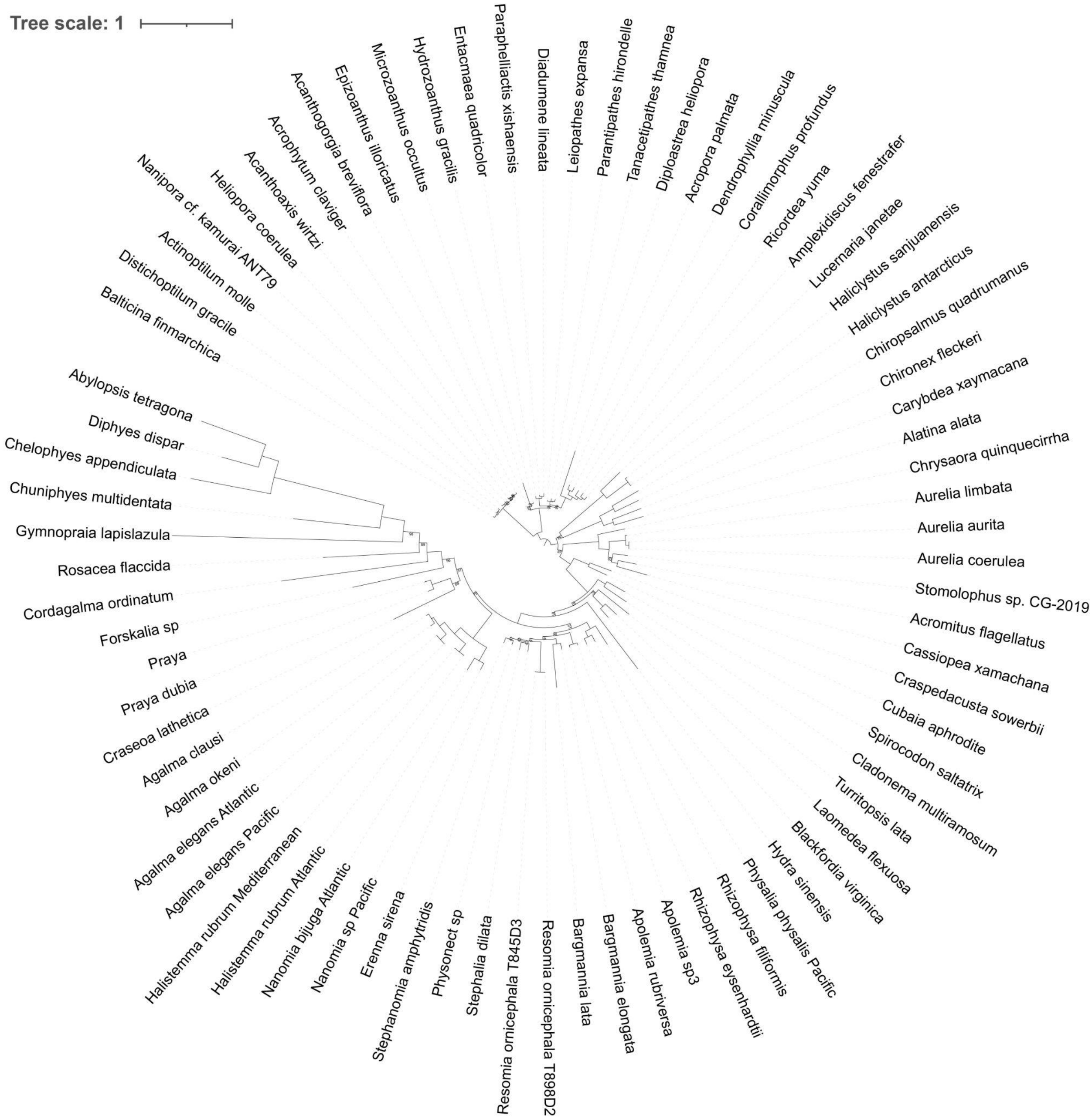


Supplementary Figure 4. Unconstrained maximum-likelihood phylogenetic tree based on 13 mitochondrial protein-coding genes. Nodes that did not show bootstrap values indicate that their bootstrap value = 100. Some nodes without maximum strength in this tree are constrained in the tree in Figure 4.

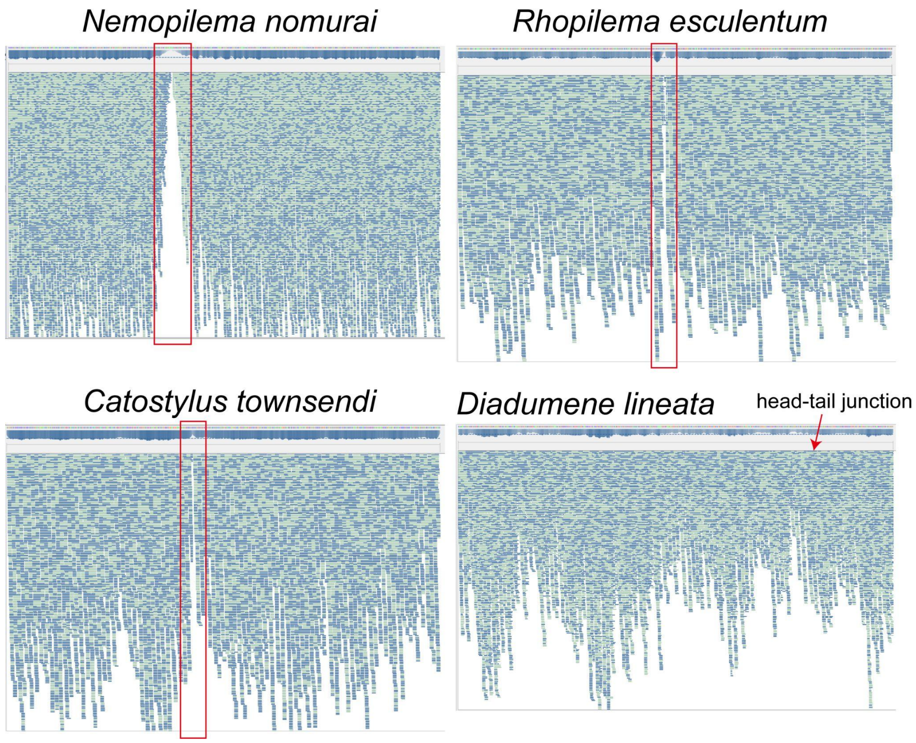


Supplementary Figure 5. Read mapping demonstrates the linear structure of *Nemopilema nomurai, Rhopilema esculentum* and *Catostylus townsendi* mitochondrial genomes. The methods used were consistent with Supplementary Figure 3. *Diadumene lineata* (an anthozoan) was a control, showing that reads pass through the head-tail junction of the circular mitochondrial genome well. Sequences and SRA data used were *Nemopilema nomurai* (KY454767, SRR6298213), *Rhopilema esculentum* (KY454768, SRR8617500), *Catostylus townsendi* (OK299144, SRR16643387) and *Diadumene lineata* (MH699974, ERR6688659).

### Supplementary tables

Supplementary tables not included in this document are available in the git repository.

Supplementary Table 1. Specimen data spreadsheet. This is the NCBI BioSample spreadsheet that was uploaded with the Illumina reads to SRA.

Supplementary Table 2. GenomeScope2 results, with additional annotations. Derived from https://github.com/dunnlab/siph_skimming/blob/main/analyses_kmer/output/genome_size_report.tsv

| **specimen** | **minimum** | **heterozygous peak** | **homozygous peak** | **manual size estimate (Gb)** | **GenomeScope size estimate (Gb)** |
|---|---|---|---|---|---|
| *Agalma elegans* Pacific | 16 | 37 | 72 | 2.44 | 2.3 |
| *Chelophyes appendiculata* | 11 | 29 | - | 1.31 | 1.2 |
| *Cordagalma ordinatum* | 8 | 22 | - | 0.74 | 0.7 |
| *Gymnopraia lapislazula* | 8 | 14 | - | 4.83 | - |
| *Nanomia bijuga* Atlantic | 17 | 44 | 89 | 0.75 | 0.7 |
| *Nanomia sp* Pacific | 10 | 19 | 37 | 1.50 | 1.4 |
| *Physalia physalis* Pacific | 15 | 30 | 58 | 1.80 | 1.7 |
| *Resomia ornicephala* T898D2 | 5 | 8 | - | 1.40 | - |

Supplementary Table 3. Manual genomes size estimates, with comparison to GenomeScope2 genome size estimates. The first three numeric values refer to the coverage of features in the histograms (Figure 3). The minimum is the position of the left-most minimum in the histogram, most k-mers to the left of this are errors. The heterozygous peak is the left-most peak (to the right of the minimum), and was used for manual genome size estimation. The homozygous peak is the second peak from the left. In some cases the homozygous counts do not form a distinct peak, but instead a shoulder to the right of the heterozygous peak. In these cases a dash is noted rather than a coverage.

Supplementary Table 4. Number of reads included in each rarefaction sample.

Supplementary Table 5. Other cnidarians mitochondrial genome information used in this study.
